# Membrane mimetics control internal hydration and catalytic activity of the copper ATPase LpCopA from *Legionella pneumophila*

**DOI:** 10.64898/2026.07.28.738645

**Authors:** Lisa Nucke, Yun-Hsuan Huang, Satoru Tsushima, Jana Oertel, Karim Fahmy

## Abstract

Membrane protein function depends critically on the surrounding lipid bilayer, which supports specific lipid–protein interactions and imposes constraints through its ensemble properties. Ion-transporting ATPases undergo large conformational changes that perturb these interactions, and the high chemical potential of water can further drive conformational transitions through changes in internal hydration. Here, we investigated the relationship between membrane environment, internal hydration, and catalytic activity in the P_1B_-type copper ATPase LpCopA from *Legionella pneumophila*. BADAN-labeled LpCopA mutants revealed distinct basal hydration states near the conserved canonical binding site (CBS) cysteines C382 and C384, with C382 residing in a more hydrated environment than C384. Using the osmolyte PEG-1500, the spectral response of BADAN indicated an average internal hydration volume of about 800 Å^3^ for LpCopA in *E. coli* lipid-doped mixed micelles (MMs), corresponding to approximately 25 water molecules. In nanodiscs (NDs), less than half of this volume responded to osmotic pressure, consistent with compaction of the transmembrane helical bundle under membrane lateral pressure. Remarkably, ATPase activity measured in different lipid reconstitution systems scaled with the extent of internal hydration, with diisobutylene/maleic acid lipid particles (DIBMALPs) imposing the tightest transmembrane compaction and lowest basal activity. These data indicate that the packing density and internal hydration of the transmembrane domain of LpCopA are strongly modulated by the membrane environment. The energetic estimates further support a model in which hydration-dependent expansion of the transmembrane domain against membrane lateral pressure contributes to the free-energy barrier of ATP hydrolysis.

## 1. Introduction

The flow of information and matter across biological membranes is vital for all cellular life. Membrane proteins acting as receptors of external signals or as transporters of ions and substrates fulfill these functions as crucial constituents of the lipid bilayers that make up the membranes of cells and organelles. The structure and function of membrane proteins is tuned by a manyfold of lipid protein interactions which stabilize an equilibrium protein structure by minimizing the free energy of the ensemble of these interactions, for example by hydrophobic matching of transmembrane protein domains to the lipid bilayer thickness, or by adaptation to bilayer curvature, lateral pressure or lipid asymmetry [1–4]. The physico-chemical linkage between membrane proteins and membrane lipids provides the ground on which the co-evolution of both can be retraced [5].

Although the lack of lipid protein interactions may lead to loss of function, membrane function itself entails structural transitions between distinct conformations which cannot share the same energy-minimized lipid protein interactions. This renders the lipidic phase an active component which affects the energetics of membrane protein function. Ion-transporting ATPases exhibit particularly large structural changes originally demonstrated for the Ca^2+^-transporting P_2_-type ATPase SERCA from the sarcoplasmic reticulum [^6^]. These transporters appear particularly well suited to study fundamental aspects of the linkage between membrane protein function and dynamic interactions within a lipid bilayer.

P-type ATPases are found in all kingdoms of life using the energy of ATP to build up and maintain ion gradients across biological membranes accompanied by transient auto-phosphorylation. In microorganisms, these transporters play a critical role in metal homeostasis and detoxification [7]. The Gram-negative pathogen *Legionella pneumophila* employs the Cu^+^-exporting P_1B_-type ATPase LpCopA in response to conditions encountered within alveolar macrophages after host infection [8, 9]. LpCopA possesses an N-terminal heavy metal binding domain and two P_1B_-type-specific transmembrane helices which precede a core structure of six helices shared by all P-type ATPases. The core structure harbors a canonical binding site (CBS) for Cu^+^ including the conserved cysteine residues C382 and C384 on helix IV, as well as other residues that define the ion pathway through the membrane domain [10]. Accessibility and affinity of these residues is under the control of three cytosolic domains found in all P-type ATPases which transmit ATP-driven structural changes to the core domain [11, 12].

For LpCopA, crystal structures have been obtained in various structural states [10, 13] and specific lipid effects on its function have been analyzed thoroughly both *in vitro* and *in* [14]. ATPase activity in mixed micelles (MMs) was found to be promoted predominantly by POPG and cardiolipin. Coarse-grained molecular dynamics simulations in lipid bilayers have traced these interactions back to defined transmembrane segments and further down to single amino acid preferences for specific lipid species [14]. However, much less is known about how the ensemble properties of the membrane environment influence internal hydration and couple to catalytic function. MMs are typically used to grow crystals and to measure ATPase function but their geometrical constraints differ markedly from those of a lipid bilayer, where lateral pressure is exerted on membrane proteins [15]. Consequently, intramolecular helix packing, hydration and hydrophobic matching constraints differ in MMs and probably also in various membrane mimetics, other than liposomes, which are increasingly used for *in vitro* catalytic assays. How these reconstitutions affect the interdependence of global bilayer properties, membrane protein hydration and catalysis has not been studied systematically.

The overwhelming chemical potential of water at the lipid protein interface favors hydrated conformations over dehydrated states. The alternating access model of the Post-Albers cycle of ion transport builds on the switch between conformations with either intra- or extracellular accessibility of an internal ion-binding site, the E1 and E2 state, respectively, with occluded states between the transition [16]. However, the analyses of E2 states of LpCopA suggested that ion release originated from ion-binding sites deep in the transmembrane domain along a water-lined pathway without passing through an occluded intermediate, emphasizing the functional importance of internal hydration [10].

We have previously shown that the CBS cysteines of LpCopA are accessible to site-directed labeling from the aqueous phase without the need for partial unfolding. This allowed us to probe the local dipolar environment near the CBS by attaching the solvatochromic fluorophore BADAN (6-bromoacetyl-2-dimethylaminonaphthalene), which revealed substantial differences in local hydration dynamics near C382 and C384 [17]. These features make LpCopA a particularly useful model system for addressing a fundamental question in membrane protein biophysics: how does the membrane environment affect internal hydration of the transmembrane domain, and how is this linked to catalytic activity?

To obtain a quantitative estimate of internal hydration, osmotic stress was applied to BADAN-labeled LpCopA mutants, allowing the number of osmotically exchangeable water molecules to be determined. This concept was originally developed for ion channels [18] and has been reviewed recently with a focus on experiments that demonstrated the functionally required water-driven expansion of the visual photoreceptor bovine rhodopsin, a prototypical class-A GPCR [19]. Such studies rely on an observable of conformational transitions that are accompanied by water uptake or release processes and, thereby, become sensitive to osmolytes. The osmotically modulated hydration processes are not required to take place at the site used for conformational monitoring as long as they are allosterically coupled to that site. In the present study, BADAN serves not only as a site-specific reporter of local polarity through its emission wavelength, but also as a probe of hydration-linked conformational equilibria through its response to osmotic stress. Here, we use these two spectral properties of BADAN to characterize both local hydration and global osmotically exchangeable water in LpCopA reconstituted in different membrane mimetics. The results show that ATPase activity correlates with internal hydration of the transmembrane domain and that this relationship is strongly modulated by the membrane environment.

## 2. Results and Discussion

### 2.1. Internal hydration of LpCopA is in osmotic equilibrium with the aqueous phase

The copper transport site of LpCopA, offers sufficient space and accessibility to allow labeling the two conserved active site cysteines C382 and C384 with BADAN in the natively folded state of the membrane protein [17]. Figure 1 shows the crystal structure of LpCopA with the approximate locations of the fluorophore when attached at either cysteine residue, together with crystallographically resolved water molecules. To enable site-specific labeling of the CBS cysteines in helix IV, the four cysteine residues in the C-terminal heavy metal-binding domain were replaced by serine.

**Figure 1.**
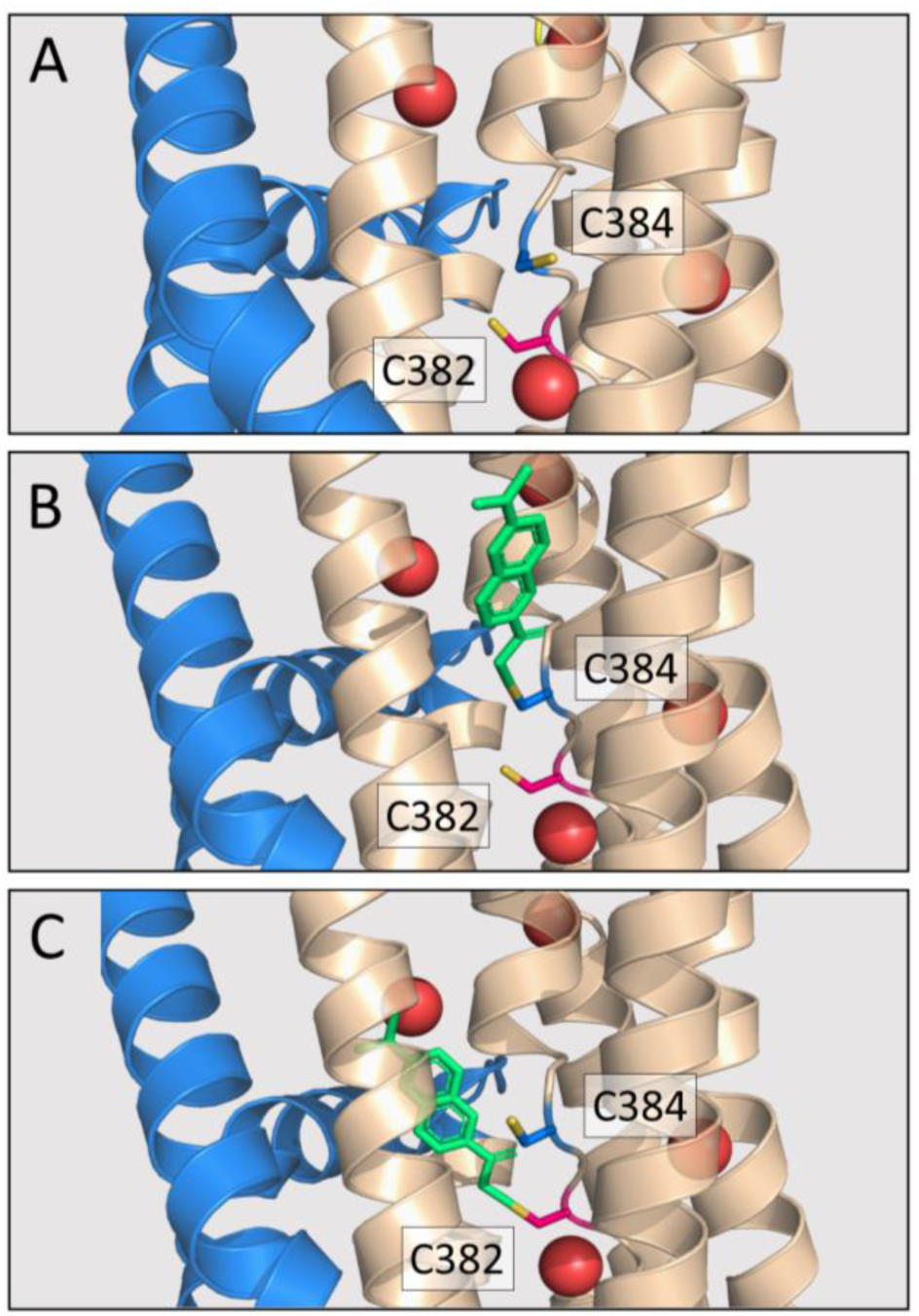
View of the conserved CPC motif in the crystal structure of LpCopA (PDB 4BBJ) with highlighted active site cysteines and water molecules. A) Wild type LpCopA B) Hypothetical orientation of the fluorophore BADAN covalently linked to cysteine 384, designated BADAN@C384. C) The equivalent model for BADAN@C382. Cysteines are in stick representation, whereas water molecules are displayed as red spheres.

We have carried out spectroscopic and functional assays of BADAN-labeled and unlabeled LpCopA to address the potential influence of the membrane environment on internal hydration and catalytic activity, respectively. In contrast to our previous time-resolved Stokes-shift experiments, which revealed different water dynamics near C382 and C384 [17], the present study was designed to estimate the total amount of osmotically exchangeable water coupled to these sites. We were particularly interested in determining whether internal hydration of the transmembrane domain depends on the membrane mimetic and whether such changes are linked to ATPase activity in the presence of *E. coli* lipids.

Figure 2 shows the effect of 32.5 % (w/w) PEG-1500 on the emission spectra of BADAN attached at either C382 or C384 in four reconstitution systems: DDM micelles, mixed micelles (MMs), nanodiscs (NDs), and DIBMALPs. The emission spectra were fit with the sum of two Gaussian components to estimate the relative populations of weakly and stronglyhydrated states near the two labeling sites. In all systems, addition of PEG-1500 shifted the spectra toward shorter wavelengths, reflected by an increased contribution of the Gaussian component with an emission maximum below 450 nm (Figure 2, green area) relative to the component above 485 nm (Figure 2, blue area). The blue shift of the BADAN emission is well-known to report the decrease of the water-dependent dipolar relaxation in the chromophore’s molecular environment in phospholipid membranes as well as in synthetic transmembrane protein segments [20, 21]. Thus, hydration near both CBS cysteines is in osmotic equilibrium with the aqueous phase and decreases upon osmotic stress.

**Figure 2.**
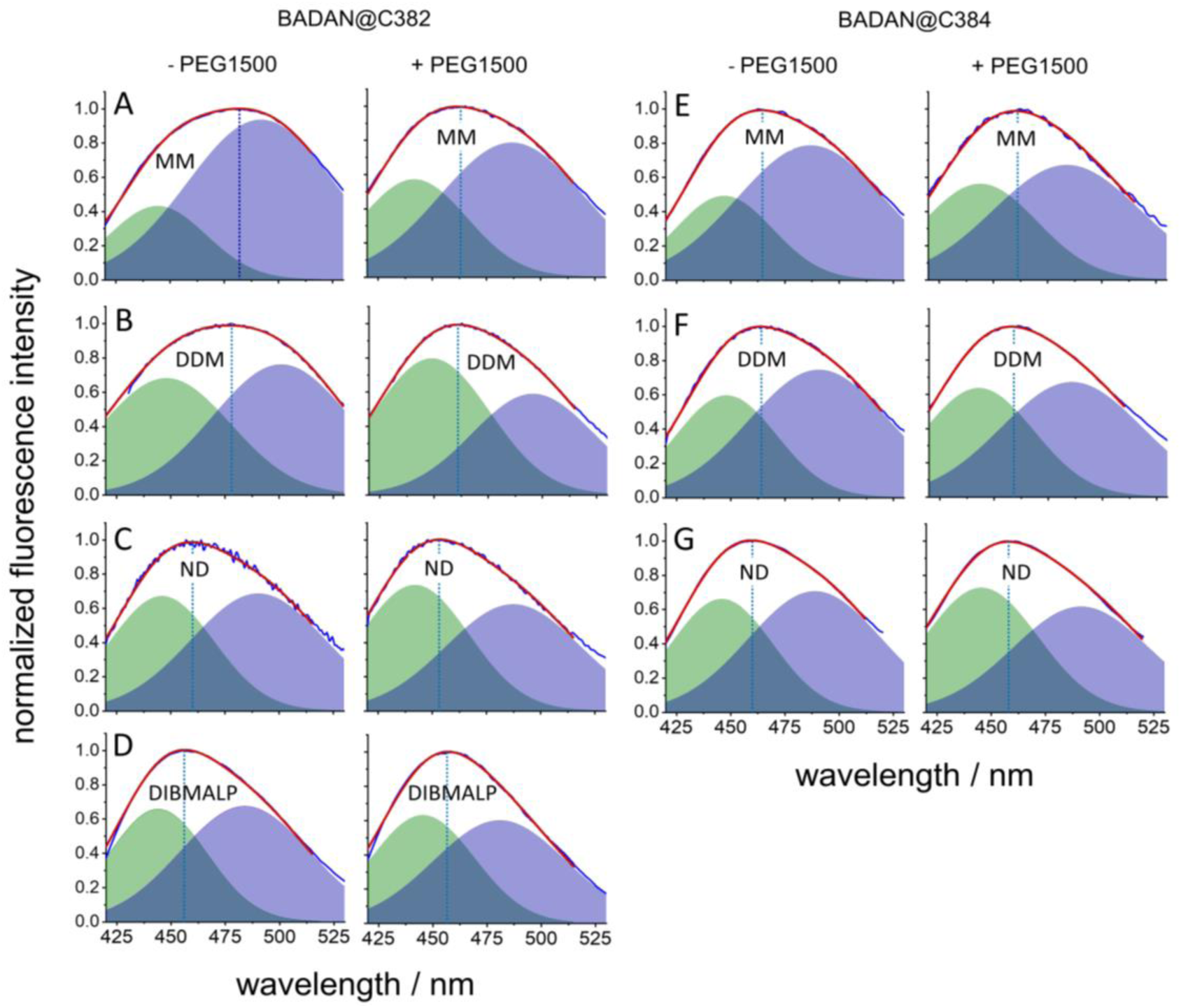
Gaussian decomposition of steady state fluorescence spectra of BADAN-labeled single-cysteine LpCopA mutants in different membrane mimetic systems with and without PEG-1500. Left panel: A-D) Fluorescence of BADAN bound to C382 in the indicated reconstitution systems, dodecyl-maltoside micelles (DDM), mixed micelles (MM), MSP1E3D1 nanodiscs (ND) and DIBMALPs. The spectra were recorded in aqueous buffer without or with 32.5 % PEG-1500 as indicated at the top of the panel. The recorded spectra (blue lines) were reproduced by the sum of two Gaussian bands (red lines) whose integrals were assigned to weakly hydrated (green areas) or strongly hydrated (blue areas) populations. Right panel: E-G) The equivalent data obtained with BADAN bound to C384.

In all reconstitution systems, basal hydration in the absence of osmolyte was higher at C382 than at C384. Moreover, the magnitude of the PEG-1500-induced spectral response followed the degree of basal hydration at the two sites, despite their separation by only a single proline residue. Compared with DDM micelles and MMs, NDs showed lower basal hydration and smaller osmolyte-induced changes. This indicates that reconstitution into NDs reduces internal hydration of LpCopA, leaving less osmotically exchangeable water within the transmembrane domain. Consistent with this interpretation, the BADAN emission spectra in NDs were more blue-shifted than in detergent-based systems even in the absence of osmolyte. This behavior agrees with our earlier observations for LpCopA reconstituted in asolectin-containing bilayers [17].

The largest osmotic modulation of internal hydration was observed in MMs rather than in pure DDM micelles. This was unexpected, because pure detergent micelles might be assumed to support the highest degree of water accessibility. By contrast, DIBMALPs exhibited the smallest osmotic response and the lowest basal hydration of all systems tested, consistent with the most compact transmembrane packing. Together, these results indicate that both the amount of internal water and its osmotic accessibility depend strongly on the membrane environment.

### 2.2. Osmotic stress reveals membrane-dependent hydration volume changes

The above qualitative description shows that osmotic sensitivity near the CBS is highly site-dependent, indicating that C382 and C384 report on distinct local hydration environments. To relate these spectral changes to the number of water molecules exchanged with the aqueous phase, we defined an apparent equilibrium constant, *K*, as the ratio of the areas of the Gaussian components assigned to the apparent strong- and weak-hydration states. The logarithm of *K* was determined for varying concentrations of PEG-1500 and plotted against the tabulated osmotic pressures of PEG-1500 solutions [22].

Figure 3 shows an approximately linear relationship between ln*K* and osmotic pressure for all reconstitution systems. This behavior indicates that, despite the simplifying two-state approximation, the spectral response can be described consistently by an apparent equilibrium of the form:

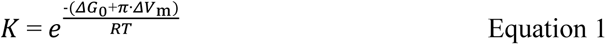

**Figure 3.**
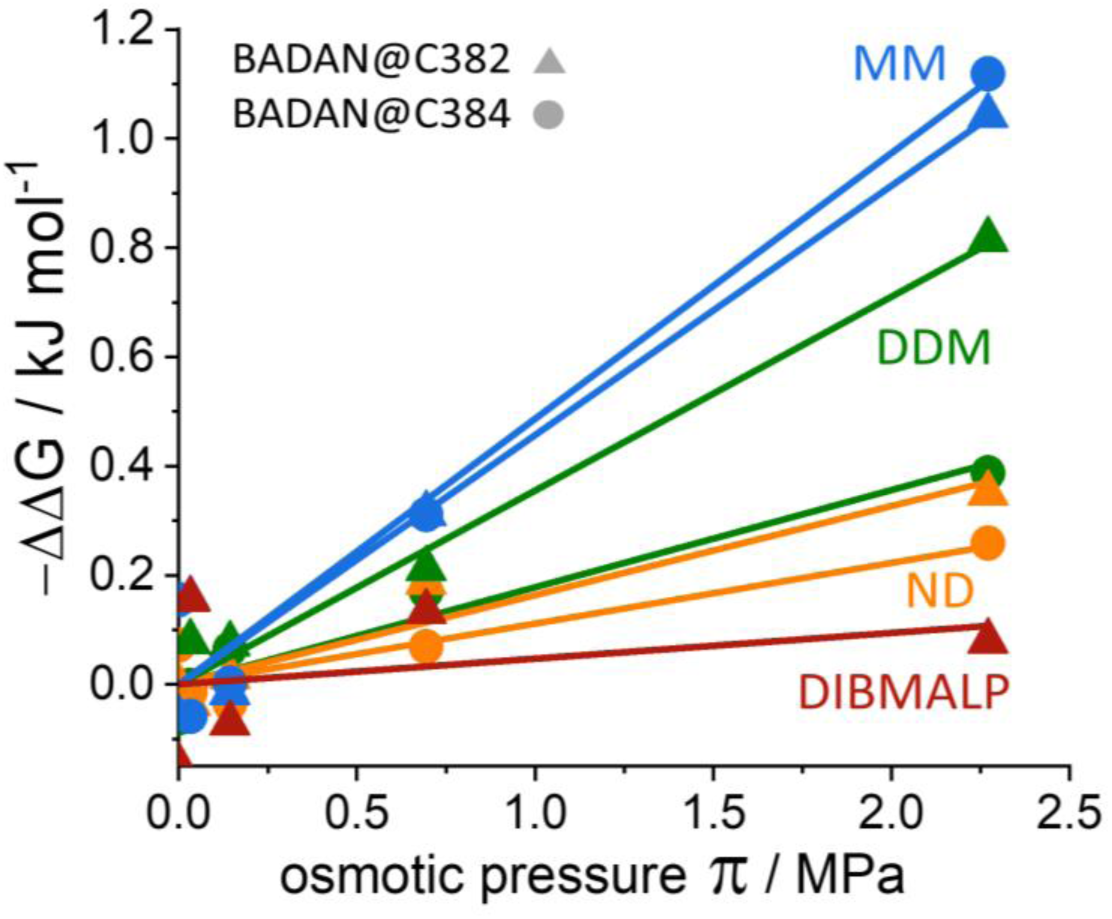
Free energy change ΔΔ*G* of the equilibrium between “strong” and “weak” hydration as a function of osmotic pressure. Emission bands of BADAN-labeled CopA single cysteine mutants (BADAN@C382, triangles; BADAN@C384, filled circles) in MMs (blue) DDM micelles (green), NDs (orange) and DIBMALPs (red) were fitted by two Gaussian bands (Figure 2). The hydration equilibrium constant was defined as the ratio of the areas under the Gaussian bands for each osmotic pressure. The slopes of the linear fits provide the value of the molar volume change -Δ*V*m upon osmotic dehydration. The derived amounts of exchanged water molecules are summarized in Table 1.

**Table 1.** Site-specific molar volume change Δ*V*m, number of exchangeable water molecules upon PEG-1500 addition, and ATPase activity of LpCopA in different reconstitution systems. The values for Δ*V*m were derived from the slope of the regression lines in Figure 3 according to Equation 1 assuming a molar water volume of 30.3 Å^3^. The labeled protein is inactivated at the active site and the unlabeled protein lacks the fluorophore for hydration monitoring. Therefore, the determination of certain parameters is not applicable (n.a.).

| Label position | Mimetic system | Emission shift / nm | $\Delta V_m / \text{nm}^3$ (error) | #H<sub>2</sub>O (error) | ATPase activity / $\mu\text{M Pi } \mu\text{g}^{-1} \text{ min}^{-1}$ (error) |
| --- | --- | --- | --- | --- | --- |
| BADAN @C384 | DDM | 464 → 459 | 0.30 (0.05) | 10.0 (1.5) | n.a. |
|  | MM | 465 → 461 | 0.81 (0.09) | 27.3 (3.2) | n.a. |
|  | ND | 460 → 458 | 0.18 (0.04) | 6.2 (1.5) | n.a. |
| BADAN @C382 | DDM | 478 → 461 | 0.60 (0.05) | 19.9 (1.7) | n.a. |
|  | MM | 482 → 463 | 0.76 (0.09) | 25.6 (3.0) | n.a. |
|  | ND | 460 → 453 | 0.27 (0.04) | 9.2 (1.5) | n.a. |
|  | DIBMALP | 456 → 456 | 0.08 (0.13) | 2.6 (4.2) | n.a. |
| No label | DDM | n.a. | n.a. | n.a. | 0.004 (0.017) |
|  | MM | n.a. | n.a. | n.a. | 0.334 (0.025) |
|  | ND | n.a. | n.a. | n.a. | 0.233 (0.033) |
|  | DIBMALP | n.a. | n.a. | n.a. | 0.070 (0.040) |

where Δ*G*_0_ is the standard free energy change of the weakly to strongly hydrated transition. The additional free energy contribution imposed by osmotic pressure is written as ΔΔ*G*_0_ = π ΔV_m_, where ΔV_m_ (> 0) is the molar volume change associated with intra-membrane protein hydration and π corresponds to the osmotic pressure. From the slope of the linear fits, we estimated Δ*V*_m_ for the different reconstitution systems from the plots of - ΔΔ*G*_0_ versus π (Figure 3). This allows approximating the total number of exchanged water molecules using 30.3 Å^3^ per water molecule (Table 1).

The data show that a total of about 25 internal water molecules is in osmotic equilibrium with the aqueous phase in MMs (Figure 3). The osmotic dehydration is equally sensed at both reporter sites, which nevertheless reside in unequal local dipolar environments as evidenced by their different emission maxima and band shapes irrespective of the presence of PEG-1500. Figures 2A and 2E show that the long wavelength integral emission is still much higher for the label at C382, than at C384 and the peak emissions from the two labeled sites stayed separated by 17 nm. In other words, the two cysteine residues still sample their differently hydrated local water pockets but may nevertheless participate in the same osmotically induced overall volume change since the slope of the regression lines obtained for both labels is virtually identical.

This is in stark contrast to the osmotic response in DDM (Figure 2B, F; Figure 3). Here, the dipolar environment of the label at C382 is linked to about 20 exchangeable water molecules, whereas only half of that value is sensed by the label at C384. In this case, two equilibrium constants with different Δ*V*_m_ are required to describe the osmotic effect. Thus, the CPC motif appears not only to separate two distinct local dipolar environments, but also to mark a region where the collective hydration behavior observed in MMs becomes partially uncoupled in DDM micelles.

LpCopA in NDs showed an intermediate behavior (Figure 2C, G; Figure 3). The amount of osmotically exchangeable water estimated from both sites was approximately half of that observed in MMs. This reduction indicates that the membrane environment in NDs favors a more compact transmembrane helical bundle with reduced internal hydration.

By comparing the effects of PEG-1500 between BADAN labeled C382 and C384, the question arises, to which extend the interrogated sites are structurally independent. They may be linked to the same osmolyte-induced global hydration changes in MMs but clearly sense different hydration sites in NDs. A structural distinction between the water accessibilities at the two sites in NDs would have functional implications. We have addressed this question by exposing LpCopA in NDs to a smaller osmolyte PEG-200. Thereby, only those internal water molecules will be extracted under osmotic stress that are connected through much smaller channels to the aqueous phase than those responding to PEG-1500 [18]. In fact, the smaller osmolyte had virtually no effect on the BADAN emission when the label was linked to C382. In contrast, PEG-200 induced a fluorescence red-shift of the label at C384 (Figure S6). Similar effects have been observed for other proteins [19] and indicate that the osmolyte enters into protein crevices, where it either interacts directly with the protein or induces internal rearrangements of hydration water without a net dehydrating effect. The different responses of the label to PEG-200 clearly demonstrate that the degree of hydration and the aqueous accessibility to C382 and C384 is shaped by the lipid environment.

The smallest hydration response was observed in DIBMALPs and was detectable only at the more strongly hydrated site, C382 (Figure 2D, Figure 3). DIBMALPs exhibited already in the absence of osmolytes the most blue-shifted BADAN fluorescence, indicating the lowest basal internal hydration of all reconstitution systems. Whereas the BADAN emission shifted from 460 nm to 453 nm in NDs (Figure 2C) upon addition of PEG-1500, the emission in DIBMALPs remained essentially unchanged at 456 nm. The modest narrowing of the emission band and the small redistribution of the underlying Gaussian components suggested only minimal dehydration. Therefore, the data evidence that the two discoidal planar lipid systems have markedly different effects on the transmembrane packing of LpCopA which leads to distinct changes in basal internal hydration as well as in the distribution width of internal dipolar micro environments of the two sampled sites.

Although the necessarily simplified analysis of static BADAN emission does not capture the full distribution of fluorescent microstates or the time-dependent decrease in photon energy during dipolar relaxation, the overall trend is clear. Relative to detergent-based systems, bilayer-based reconstitution in NDs and especially in DIBMALPs reduces internal hydration of the transmembrane domain. For both BADAN-labeled mutants, transfer from detergent to NDs caused an approximately two-fold decrease in osmotically exchangeable water while preserving the higher basal hydration at C382 compared with C384. DIBMALPs represent the limiting case, in which osmotically exchangeable water was nearly absent and the BADAN emission at the CBS was maximally blue-shifted. These observations are consistent with a more tightly packed and less hydration-permissive transmembrane helical bundle in DIBMALPs.

### 2.3. Internal hydration correlates with catalytic activity

The pronounced differences in internal hydration among the reconstitution systems prompted us to ask whether these differences correlate with ATPase activity. Figure 4 compares the ATPase activity of unlabeled, wild-type LpCopA in MMs, DDM, NDs and DIBMALPs, measured as the time-dependent release of inorganic phosphate in the presence of ATP and Ag^+^ as a substitute for Cu^+^. The observed ATPase activities were generally comparable to previously reported values [14], with activity highest in MMs and lower in NDs. By contrast, LpCopA was almost inactive in pure detergent and DIBMALPs.

**Figure 4.**
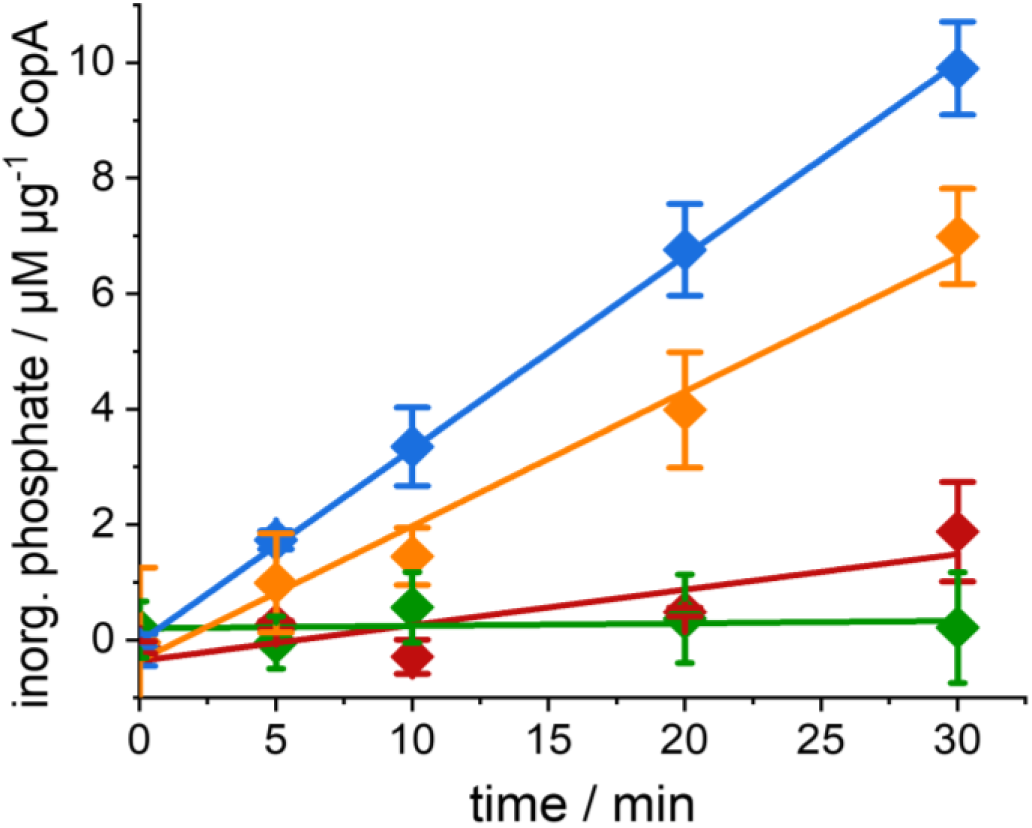
Time-dependent release of inorganic phosphate during the catalytic cycle of LpCopA in MMs (blue), NDs (orange), DDM micelles (green) and DIBMALPs (red). ATP and Ag^+^ were 5 mM and 500 µM, respectively, in a final reaction volume of 160 µL. Experiments have been performed in duplicates. The catalytic rates were obtained from the slope of the regression lines as summarized in Table 1.

Figure 5 compares the osmotic response of the BADAN label at C382 in the different reconstitution systems with the ATPase activity of the unlabeled protein in the same membrane mimetic. The comparison reveals a correlation between catalytic activity and the amount of osmotically exchangeable water for all reconstitution systems, except in pure DDM. Thus, internal hydration of the transmembrane domain appears to be functionally important, but its extent is strongly dependent on the membrane environment. Functional impairment of membrane proteins in pure detergent has been described extensively before [14, 23–25] and is commonly attributed to the loss of essential protein-lipid interactions. In contrast, the lack of catalytic activity in DIBMALPs suggests that the DIBMALP environment imposes a level of transmembrane compaction that restricts the hydration changes required for catalytic turnover. Whether the low catalytic turnover in DIBMALPs reflects an over-constrained *in vitro* system or approximates the tightly regulated basal state of LpCopA under native membrane lateral pressure remains to be established.

**Figure 5.**
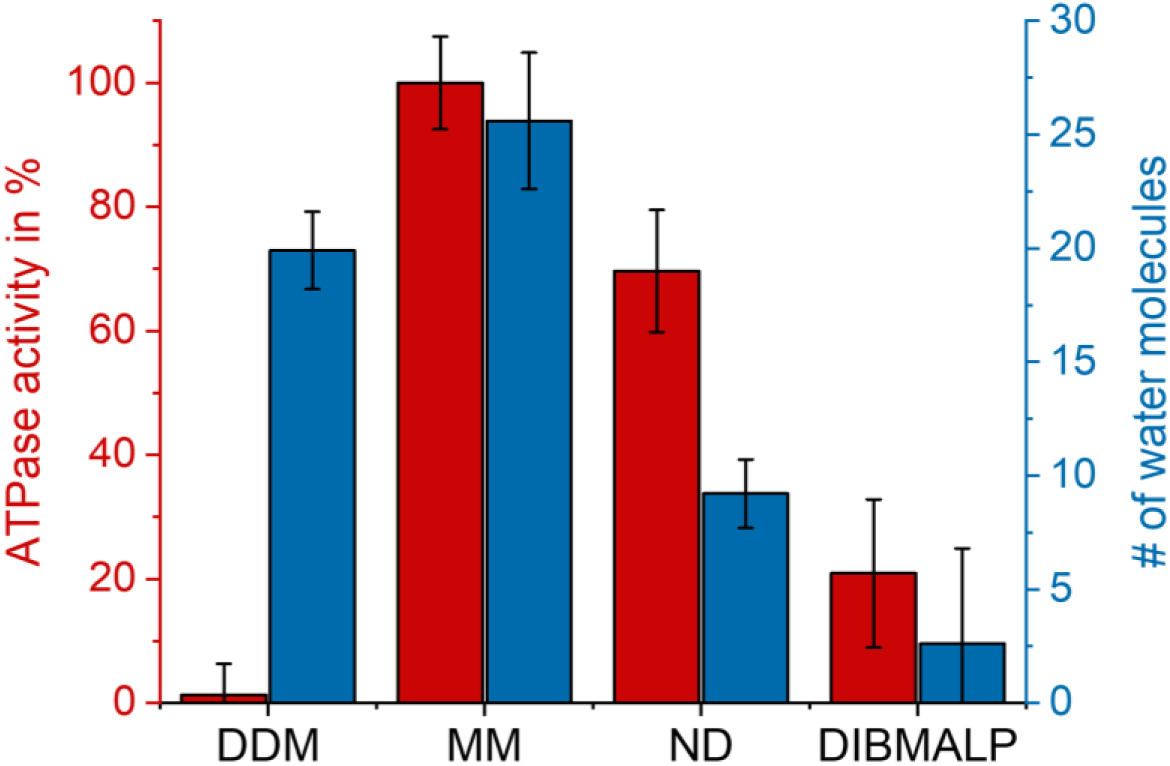
Comparison of LpCopAs ATPase activity and hydration properties of the active site cysteine 382 in different membrane mimetic systems. The ATPase activity of WT CopA as well as the site specific hydration change of the BADAN@C382 mutant has been determined in the following membrane mimetic systems: dodecyl maltoside micelle (DDM), mixed micelle (MM), MSP1E3D1 nanodisc (ND) and DIBMA (DIBMALP). 100% activity corresponds to a value of 0.334 ± 0.025 µM Pi µg^-1^ min^-1^. Further values for ATPase activity and number of water molecules can be found in Table 1.

### 2.4. Mechanistic aspects of hydration-dependent catalytic activity of LpCopA

The dependence of ATPase activity on the reconstitution system has direct mechanistic implications, particularly when comparing MMs and NDs. The rate of ATP hydrolysis was about 43 % higher in MMs than in NDs, indicating that a planar bilayer imposes higher energy barriers along the trajectory of functionally required conformational changes of LpCopA than the much less constraint, flexible MMs. We hypothesize that energy barriers result from the transient hydration-driven expansion of LpCopA against membrane lateral pressure in NDs. This is in line with the reduction of internal hydration of LpCopA by either osmotic stress on the MM or by reconstitution into NDs which themselves exert a dehydrating effect upon compaction of the transmembrane helical bundle.

In order to estimate the apparent free energy difference Δ*G*_cmp_ for compacting the LpCopA structure from the more hydrated MM state to the less hydrated ND state, we compared the internal hydration equilibria in the absence of osmolytes. The Gaussian band intensity ratios of 0.36 in MMs and 0.84 in NDs for BADAN@C382 (Figure 6) correspond to an apparent free energy difference of approximately 2 kJ mol^-1^ between the more hydrated MM state and the less hydrated ND state. Independently, the osmolyte experiments indicated that the MM and ND states differ by an osmotically exchangeable volume of approximately 500 Å^3^ at the C382 reporter site (Table 1). Assuming a bilayer thickness of 5 nm and approximating the transmembrane helical bundle as a cylindrical constriction, this volume difference corresponds to an effective cross-sectional area change Δβ_cs_ of approximately 10 Å^2^, which entails work of compaction against lateral pressure. This simplified model cannot capture the full lateral pressure profile [15], but provides a plausible approximation of the mechanical energy of compaction Δ*G*_cmp_ in the form of:

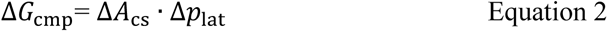

**Figure 6:**
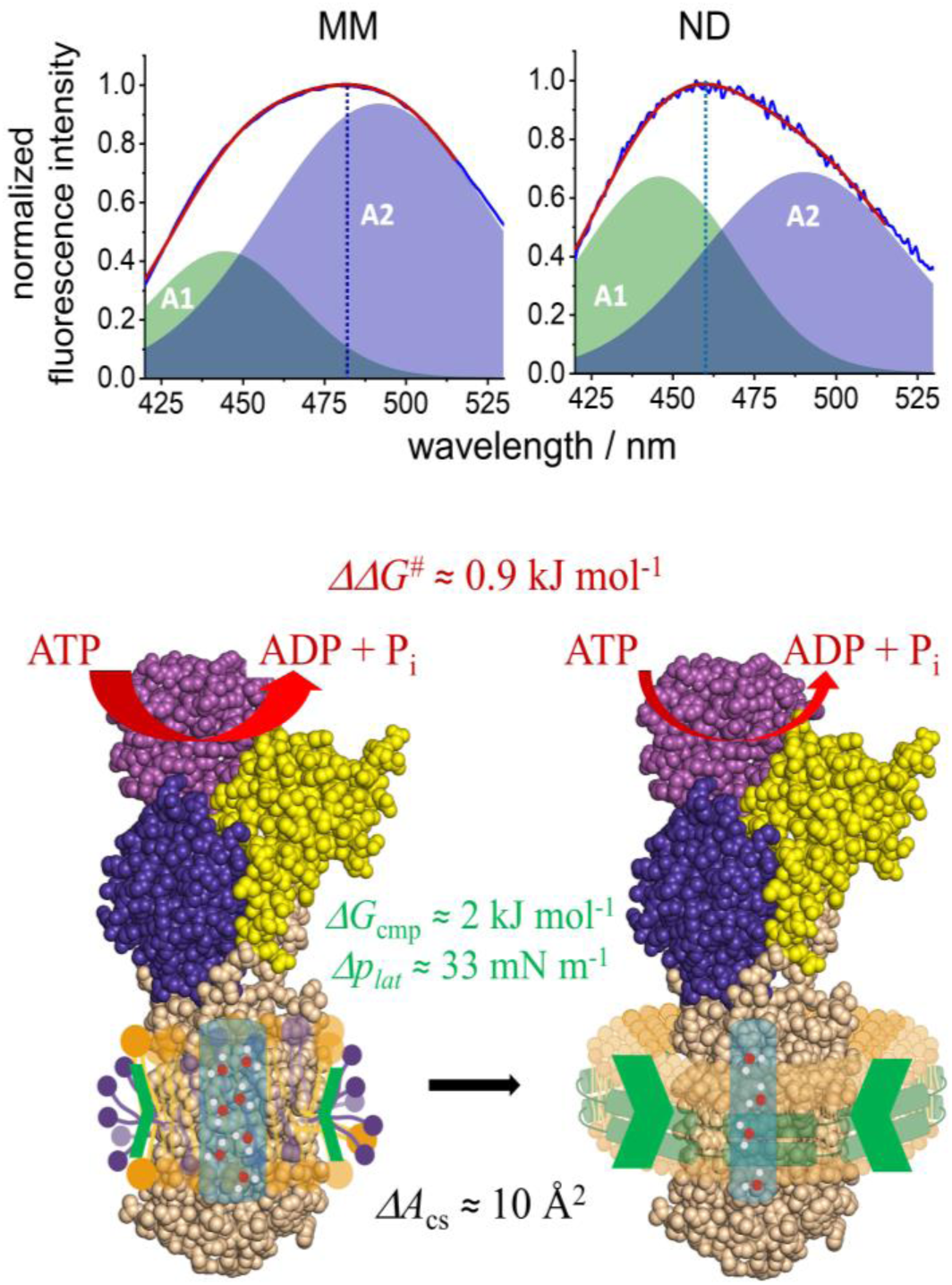
Estimates of the energetic and geometric differences between LpCopA reconstituted in a mixed micelle (MM) and a MSP1E3D1 nanodisc (ND) which provides a bilayer environment. The Δ*G*cmp between MMs and NDs has been derived from the change of the strongly hydrated (A2, blue area) to the weakly hydrated (A1, green area) population ratio in the emission spectra of BADAN linked to C382 (upper panel). The lateral pressure difference Δ*p*lat between the membrane mimetics was calculated from Δ*G*cmp for a 5 nm long cylindrical volume. ΔΔ*G*^#^ was directly obtained from the ratio of catalytic rates in the two reconstitutions (Figure 4; Table 1).

where Δ*p*_lat_ denotes the difference in bilayer-imposed lateral pressure between MMs and NDs. Attributing the entire apparent free-energy difference of approximately 2 kJ mol^-1^ to this contribution yields a lateral-pressure equivalent of 33 mN m^-1^ upon transfer of LpCopA from MMs into NDs. However, the transition from MMs to NDs also changes membrane curvature, lipid-chain ordering and bilayer thickness. Consequently, Δ*G*_cmp_ may include contributions from curvature-dependent deformation, hydrophobic mismatching, and other membrane–protein interactions that are not explicitly represented by Equation 2. The value of 33 mN m−1 should therefore be rather regarded as a mechanical estimate. Nevertheless, its magnitude is comparable to the equivalence surface pressure between lipid monolayers and bilayers of 35 mN m^-1^ [26].

In an analogous way, the difference in catalytic rates between the two reconstitutions can be expressed as an apparent activation free energy difference of ΔΔ*G^#^* ≈ 0.9 kJ mol^-1^ (Table 1). If this kinetic difference is interpreted within the same simplified mechanical framework introduced above, and using the effective stress scale of Δ*p*_lat_ ≈ 33 mN m^-1^, it corresponds to a cross-sectional area change Δ*A*_cs_ of approximately 4.5 Å^2^. This value is smaller than, but of the same order of magnitude as, the area change estimated from the osmolyte-derived hydration volume difference between MMs and NDs. Thus, both analyses are consistent with the notion that relatively small hydration-linked changes in transmembrane packing can accompany the different functional states supported by the two membrane mimetics. Figure 6 summarizes the derived energetic and geometrical quantities.

Taken together, the osmolyte experiments revealed the existence of exchangeable intra-membrane protein water in LpCopA. Hydration was not uniform around the canonical binding site, as C382 and C384 differed in their basal environments and, depending on the membrane mimetic, also in their coupling to osmotic stress. The total amount of exchangeable water varied strongly between reconstitution systems and was highest in mixed micelles, reduced in nanodiscs, and nearly absent in DIBMALPs. These differences paralleled the catalytic behavior of the unlabeled protein, with nanodiscs supporting partial activity whereas DIBMALPs did not. The energetic estimates further support the interpretation that membrane-dependent compaction of the transmembrane domain, and the associated reduction in internal hydration, contribute to the free-energy cost of catalysis.

## 3. Conclusions

We have shown that different membrane mimetics of identical lipid composition can nevertheless strongly modulate the hydration state of the transmembrane domain of LpCopA and affect its catalytic activity. Mixed micelles supported the largest amount of osmotically exchangeable internal water and the highest ATPase activity, nanodiscs exhibited intermediate levels of both, whereas DIBMALPs imposed the tightest compaction, with correspondingly minimal osmotic responsiveness and catalytic turnover.

The data suggest that changes of internal hydration during the Post-Albers cycle are required for catalytic activity. Using the experimentally derived lateral pressure in NDs and a simplified geometrical model, we hypothesize that protein expansion by a few water molecules contributes to the activation energy of the ATPase. In this view, internal hydration is not merely a passive structural feature but a functionally and kinetically important part of the energetic landscape of the transport cycle of LpCopA.

Future work will be required to resolve how these hydration changes map onto specific conformational states of the catalytic cycle and how they relate to ion binding, release, and transmembrane rearrangement.

## 4. Experimental Section

### Overproduction and purification of LpCopA

Wild-type LpCopA as well as LpCopA mutants (single-cysteine mutants BADAN@C382 and BADAN@C384, and the cysteine-free mutant, cf) were produced and purified as previously described [17]. Additional details are provided in the Supporting Information.

### Site-directed labeling of LpCopA

To obtain site-specific information on the polarity of the active-site microenvironment, the fluorescence probe 6-bromoacetyl-2-dimethylaminonaphthalene (BADAN) was covalently attached to either cysteine residue of the CPC motif on helix M4. Nonspecific labeling was minimized by employing an on-column labeling strategy. Protein expression, cell lysis, solubilization, and centrifugation were performed as described in the section *Overproduction and purification of LpCopA* (see **Supporting Information**).

The sample was adjusted to 50 mM imidazole and loaded onto a 1mL Ni-NTA HisTrap column equilibrated with KT buffer (50 mM Tris, pH 7.4, 200 mM KCl, 1 mM MgCl₂, and 10 % glycerol) supplemented with 0.25 % (w/v) DDM and 50 mM imidazole. Bound protein was labeled by washing with 15 column volumes (CVs) of BADAN wash buffer (50 mM imidazole, 0.25 % (w/v) DDM, 15 µM BADAN dissolved in KT buffer) at a flow rate of 0.3 mLmin⁻¹. Labeling was followed by extensive washing with 70 CVs of equilibration buffer and 20 CVs of KT buffer containing 0.15 % (w/v) DDM and 50 mM imidazole. The protein was eluted using a linear imidazole gradient from 50 to 500 mM. PD-10 desalting columns (Cytiva) were used to exchange the labeled protein into buffers suitable for downstream applications.

### Overproduction and purification of MSP1E3D1

The plasmid pMSP1E3D1 was obtained from Addgene (plasmid #20066). Details on overproduction and purification of MSP1E3D1 are provided in the Supporting Information.

### Preparation of DIBMA

DIBMA polymer powder (Acusol 460ND, kindly provided by the group of Prof. Sandro Keller) was dissolved in water to a final concentration of 200 mg mL⁻¹. Dissolution was facilitated by alternating incubation in a 60 °C water bath and vortexing. Three milliliters of the DIBMA solution were dialyzed against 1 L of buffer A (50 mM Tris, pH 7.4, 200 mM NaCl) using a 3.5 kDa MWCO dialysis membrane (SpectraPor) at room temperature for a total of 16 h, with buffer replacement after 8 h. After dialysis, the solution was filtered through a 0.22 µm filter. To enable accurate determination of membrane-to-polymer mass ratios, the dialyzed DIBMA solution was lyophilized using a SpeedVac (Thermo Fisher). The resulting DIBMA powder was stored at 4 °C.

### Assembly and purification of LpCopA in MSP1E3D1 nanodiscs

LpCopA or its mutants were exchanged into HNMS buffer (50 mM HEPES, pH 7.4, 200 mM Na₂SO₄, 5 mM MgSO₄) containing 0.1 % (w/v) DDM and 3 mM BME using PD-10 columns and subsequently concentrated using Macrosep Advance centrifugal devices (30 kDa MWCO, Pall Corporation). E. coli polar lipid extract (Avanti) was dissolved to a concentration of 4 mM in water containing 12 mM DDM. The lipid solution was vortexed and sonicated until homogeneous.

LpCopA, MSP1E3D1, and lipids were mixed at a molar ratio of 1:6:200 (3–7 µM CopA) and incubated on ice for 1 h. Detergent removal was performed using detergent removal spin columns (Pierce). The flow-through was adjusted to 20 mM imidazole and applied to a HisTrap spin column (GE Healthcare) to separate empty lipid nanodiscs from protein-containing nanodiscs. Elution fractions were pooled, centrifuged at 20,000 × g for 10 min at 4 °C, and further purified by size-exclusion chromatography using a Superdex 200 Increase 10/300 GL column (GE Healthcare) to obtain monodisperse nanodiscs (Figure S1).

### Formation of CopA filled DIBMA lipid particles

DIBMA enables direct solubilization of native membranes together with their intrinsic membrane proteins [27]. Accordingly, DIBMA was added to membranes isolated from CopA-overproducing *E. coli* strains. Cell culture, lysis, and membrane extraction were performed as described above, except that the lysis buffer contained 50 mM Tris and 200 mM NaCl supplemented with protease inhibitors, DNase I, and lysozyme. The membrane pellet was weighed, disrupted using a Dounce homogenizer, and adjusted to a concentration of 60 mg mL⁻¹ with lysis buffer.

DIBMA was added at a mass ratio of 0.83 (*m*_DIBMA_/*m*_wet membrane_), and the mixture was incubated for 2 h at room temperature with agitation. Insoluble material was removed by ultracentrifugation at 100,000 × g for 1 h. In parallel, 2 mL of PureCube Ni-NTA agarose (Cube Biotech) were equilibrated in a spin column with buffer containing 50 mM Tris, 200 mM NaCl, 10 mM imidazole, and 3 mM BME. The solubilized sample was incubated with the resin overnight at room temperature on an end-over-end shaker. After batch binding, the flow-through was collected, followed by four wash steps (50 mM Tris, 200 mM NaCl, 20 mM imidazole, 3 mM BME). His-tagged LpCopA DIBMALPs were eluted stepwise using increasing imidazole concentrations (250, 300, 450, and 500 mM; 2 mL each). PD-10 columns were used for imidazole removal. To eliminate aggregates, the eluate was centrifuged at 100,000 × g for 1 h at 4 °C (Figures S2 and S3).

### BADAN labeling of LpCopA in E.coli membranes prior to DIBMA solubilization

Because direct formation of DIBMALPs precludes on-column labeling of detergent-solubilized LpCopA, BADAN was added directly to *E. coli* membranes containing overexpressed LpCopA. The membrane harvesting step was performed as described above. This was followed by extensive washing and dilution with additional *E. coli* lipids prior to DIBMA-mediated solubilization. Cell culture, lysis, and membrane extraction were carried out as described above. Membranes were resuspended using a Dounce homogenizer, and BADAN dissolved in DMF was added to a final concentration of 30 µM. The membrane-BADAN mixture was incubated for 30 min at 4 °C.

Membranes were pelleted by ultracentrifugation at 100,000 × g for 1 h at 4 °C, resuspended in buffer by Dounce homogenization, and centrifuged again. This wash cycle was repeated four times to promote partitioning of unbound BADAN into the aqueous phase. The wet membrane mass was determined, and an equal mass of *E. coli* membranes lacking CopA was added. The combined membranes were solubilized with DIBMA at a mass ratio of 0.83 (*m*_DIBMA_/*m*_wet membrane_). Further processing of BADAN labeled DIBMALPs was performed as described in the methods part *Formation of CopA filled DIBMA lipid particles*.

### ATPase assay and malachite green assay

During copper transport, ATP is hydrolyzed to ADP and inorganic phosphate. The amount of released phosphate was quantified using a colorimetric malachite green assay to assess ATPase activity [27–29]. Assay conditions were largely adapted from previous work [14]. CopA samples in different membrane-mimetic systems were exchanged into ATPase assay buffer (50 mM HEPES, pH 7.0, 75 mM Na₂SO₄, 2 mM TCEP, and 5 mM MgSO₄) using PD-10 columns. For lipid–detergent micelles, 0.28 mM C₁₂E₈ was included in the assay buffer, and 0.14 mg mL⁻¹ solubilized *E. coli* lipids were added directly to the protein solution.

AgNO₃ was added to a final concentration of 500 µM, and samples were incubated at 37 °C. Reactions were initiated by addition of 5 mM ATP. At defined time points (0, 5, 10, 20, and 30 min), aliquots (160 µL) were quenched by rapid freezing in liquid nitrogen. Phosphate quantification was based on formation of a phosphomolybdate complex, followed by malachite green intercalation. Strict timing of pipetting steps was essential: at *t* = 0 min, 50 µL of 20 % trichloroacetic acid were added, samples were thawed at 70 °C for 15 s, and vortexed. At *t* = 2 min, samples were centrifuged for 1 min at 13,200 rpm. At *t* = 3.5 min, 60 µL of supernatant were transferred into tubes containing 100 µL acidic molybdate solution. At *t* = 3.75 min, 30 µL malachite green solution were added. At *t* = 5.75 min, free malachite green was degraded by addition of 200 µL 7.8 % H₂SO₄. After 70–100 min of color development, 200 µL were transferred to a 96-well plate and absorbance was measured at 625 nm.

### Sample preparation for osmolyte experiments

Stock solutions of 50 % (w/v) PEG were prepared in 50 mM HEPES, 75 mM Na₂SO₄, 5 mM KNO₃, 5 mM MgSO₄, and 2 mM TCEP at pH 7.0 and filtered through 0.22 µm sterile filters. Stock solutions were mixed with protein samples to yield final osmolyte concentrations between 2.5 % and 32.5 %. Samples were equilibrated for 15 min at 20 °C under constant agitation prior to fluorescence measurements. Osmotic pressure was calculated for the corresponding PEG concentrations using the polynomial expression [22]:

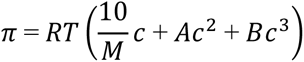

where *M* is the molecular mass of the osmolyte and *c* (g dL⁻¹) is the osmolyte concentration. The second and third virial coefficients for PEG1500 were taken as *A* = 4.7 × 10⁻⁴ and *B* = 0.9 × 10⁻⁵, with units defined as *A* = *a*(10/*M*)² and *B* = *b*(10/*M*)³ [22]. PEG concentrations were converted from g per 100 g to g dL⁻¹ using the density δ of PEG solutions at 298 K, which depends on the mass fraction *w* (g g⁻¹) but not on PEG molecular mass [30]:

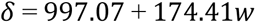

### Fluorescence measurements

Steady-state fluorescence spectra of LpCopA in different osmolyte solutions were recorded using an LS 55 luminescence spectrometer (PerkinElmer). Excitation was set to 385 nm with a bandwidth of 5 nm for both excitation and emission. Spectra were collected at a scan speed of 100 nm min⁻¹.

## Acknowledgements

We acknowledge helpful scientific discussions with Ahmed Sayed and technical assistance by Jenny Philipp (Helmholtz-Zentrum Dresden-Rossendorf). Expert support by Thomas Kurth from the electron microscopy facility (Technische Universität Dresden) is gratefully acknowledged, as well as IT support by Ronny Berndt (Helmholtz-Zentrum Dresden-Rossendorf). Furthermore, we thank Prof. Sandro Keller (Universität Graz) for kindly providing Acusol 460ND.

