## Supplemental Material for "Membrane mimetics control internal hydration and catalytic activity of the copper ATPase LpCopA from *Legionella pneumophila*"

### Supplementary Experimental Section

#### Overproduction and purification of LpCopA

Overproduction and purification protocols of wild-type LpCopA followed previously described procedures [1]. In short, the membrane protein has been overproduced in *E. coli* C43(DE3). Therefore, 10 ml LB medium + 100 µg ml$^{-1}$ Ampicillin starter culture were incubated over night at 37 °C and 180 rpm. On the next day, 500 ml LB medium + 100 µg ml$^{-1}$ Ampicillin were inoculated with 2.5 ml starter culture and incubated at 37 °C, 180 rpm until OD 0.7 has been reached. To induce gene expression 0.5 mM IPTG were added to the culture medium and cells were incubated at 18 °C for 4 h. The cell pellet was obtained by centrifugation at 6000 x g, 4 °C for 10 min, resuspended in PBS buffer, transferred to 50 ml Falcon tubes and again centrifuged. Prior to lysis, *E. coli* cells were resuspended in KT buffer (50 mM Tris, pH 7.4, 200 mM KCl, 1 mM $MgCl_2$ and 10 % glycerol) adding freshly 3 mM β-mercaptoethanol (BME), protease inhibitor and 12 µg ml$^{-1}$ DNaseI. Three passages through the high pressure homogenizer at 15,000 psi ruptured the cells. Cell debris has been removed by centrifugation at 10,000 x g, 4 °C for 30 min, and solid detergent of a final concentration of 0.8 % (w/v) has been added to the supernatant. The solubilization process took place for 40 min at 4 °C under constant agitation.

Afterwards, removal of unsolubilized material has been done at 100,000 x g, 4 °C for 40 min. 50 mM imidazole were added to the supernatant which has been loaded onto a 1 ml Ni-NTA HiTrap column equilibrated with KT buffer supplemented with 3 mM BME, 50 mM imidazole and 0.25 % (w/v) detergent (equilibration buffer). After washing with 15 CVs equilibration buffer and 2 CVs equilibration buffer containing 0.15 % (w/v) detergent, the protein was eluted with KT buffer, 3 mM BME, 0.15 % (w/v) detergent and 500 mM imidazole. Collected elution fractions were directly desalted using PD10 columns into KT-buffer, 3 mM BME, 0.15 % (w/v) detergent and 20 % glycerol. Proteins were frozen in liquid nitrogen and stored at -70 °C. For cysteine mutants (BADAN@C382, BADAN@C384 and cf), BME has not been added to the buffers.

**Overproduction and purification of MSP1E3D1**

The plasmid pMSP1E3D1 was obtained by Addgene (plasmid #20066) and has been used to overproduce and purify MSP1E3D1 in *E. coli* BL21(DE3) as described previously [1-3]. 30 µg $ml^{-1}$ kanamycin were supplemented to the double-strength YT medium and cells were grown to an OD 600 of 2.5 – 3 prior to induction with 0.3 mM IPTG. Harvested cells were resuspended in lysis buffer (50 mM Tris, pH 7.4, 200 mM NaCl) supplemented with protease inhibitors and 12 µg $ml^{-1}$ DNaseI. Upon cell lysis through a high pressure homogenizer (1x 500 psi, 3x 15,000 psi) and removal of aggregates by centrifugation, the collected supernatant supplemented with 28 mM imidazole has been loaded onto a prepacked His-Trap HP column. Immobilized metal ion affinity deviates from previous protocols by an additional wash step with cholate (50 mM Tris, pH 7.4, 200 mM NaCl, 1 % (w/v) cholate). The His-tag was cleaved off with His-tagged TEV protease (200:1 (w/w), 20 h, room temperature) and the protease was removed on a His-Trap HP column (GE healthcare). MSP1E3D1 was concentrated using Vivaspin15 Turbo filters with 10 kDa MWCO, flash frozen and stored at -70°C.

**Protein quantification**

Quantitative protein quantification has been performed using either UV-Vis spectroscopy (Instruments: NanoDrop 2000c, Nanodrop Technologies or Lambda 35 spectrophotometer, Perkin Elmer) or a Lowry based, detergent compatible protein assay (Bio-Rad Laboratories, Inc.). Concentration calculations with absorption at 280 nm values were performed with the following molar extinction coefficients: 65890 $M^{-1}$ $cm^{-1}$ for CopA in detergent or mixed micelles and 119750 $M^{-1}$ $cm^{-1}$ for CopA in MSP1E3D1 nanodiscs.

##### Transmission electron microscopy

In order to evaluate size and homogeneity of LpCopA in DIBMA nanodiscs, they were monitored by transmission electron microscopy (TEM) as follows: 5-10 µl protein solution were adsorbed to a 300 carbon mesh for 10 min at room temperature. The grid has been washed twice with a drop of water, and was negatively stained with 1 % (w/v) uranyl acetate for 20 s. Excess stain has been removed with filter paper. Images were collected with a Jeol Ruby CCD camera on a 120 kV Jeol JEM 1400Plus transmission electron microscope. Measurements were performed by Dr. Thomas Kurth (CMCB-electron microscopy facility).

##### Dynamic light scattering

Dynamic light scattering was performed on a Zetasizer Nano ZS (Malvern Instruments) equipped with a 632.8 nm laser. The 173° backward scattering has been monitored by the photodetector. Samples were measured in a low-volume quartz batch cuvette with 3 mm pathlength (Hellma Analytics). Attenuator settings were automatically optimized by the instrument.

### Supplementary Figures

#### Reconstitution of LpCopA nanodiscs

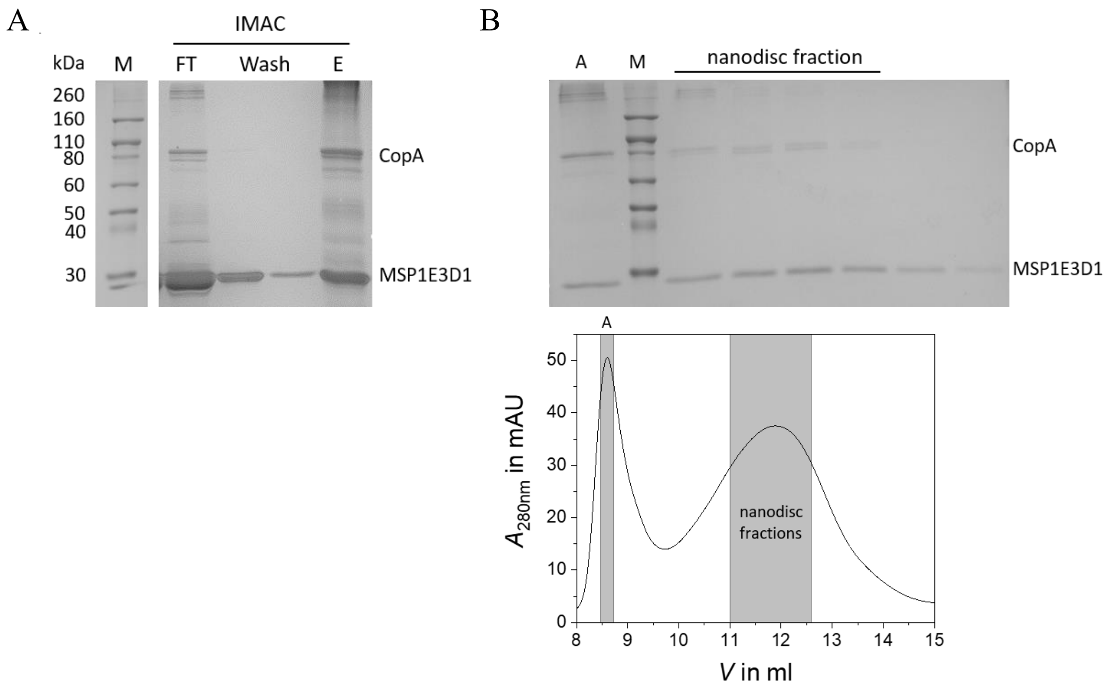

**Figure S1.** Reconstitution of LpCopA into MSP1E3D1 nanodiscs. **(A)** SDS-PAGE analysis of IMAC separating empty and filled LpCopA nanodiscs. M: Novex Sharp Prestained protein size standard, FT: Flow through, Wash, Wash fractions and E: Eluted fractions. **(B)** SEC chromatogram of IMAC eluate fraction (lower panel) and SDS-PAGE analysis of collected SEC fractions (upper panel). Two prominent peaks were observed in the size exclusion chromatogram. A - Elution at the void volume of 8.5 ml and hence corresponding putatively to aggregated particles. Nanodisc fraction - Elution at 11.9 ml and corresponding to LpCopA MSP1E3D1 nanodiscs, which could be verified by the LpCopA and MSP1E3D1 protein band on the SDS PAGE gel.

#### Preparation of LpCopA DIBMALPs

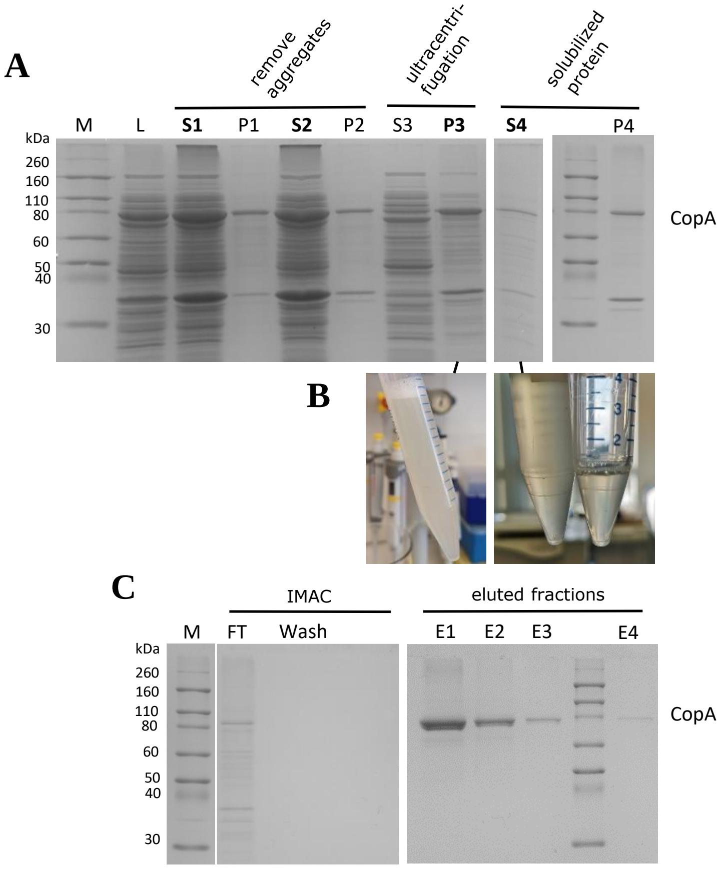

**Figure S2.** Preparation of LpCopA DIBMALPs. **(A)** SDS-PAGE analysis of different preparatory steps during LpCopA-DIBMALP production. M: Novex Sharp Prestained protein size standard. L: Cell lysate. $SN_{1\text{-}2}$ and $P_{1\text{-}2}$: Supernatant and pellet after centrifugation to remove aggregates. $SN_3$ and $P_3$: Supernatant and pellet after ultracentrifugation to harvest membrane fraction. $SN_4$ and $P_4$: Supernatant and pellet after solubilization of membrane fraction by DIBMA and ultracentrifugation to remove unsolubilized material. **(B)** Sample of harvested membrane fraction before (left) and after (right) the addition of DIBMA. **(C)** SDS-PAGE analysis of IMAC to separate LpCopA filled DIBMALPs from empty and other protein-DIBMALPs. FT: Flow through of IMAC. Wash: Wash fraction of IMAC. $E_{1\text{–}4}$: Collected elution fraction of IMAC after buffer exchange with PD10 columns containing LpCopA DIBMALPs.

#### Characterization of LpCopA DIBMALPs

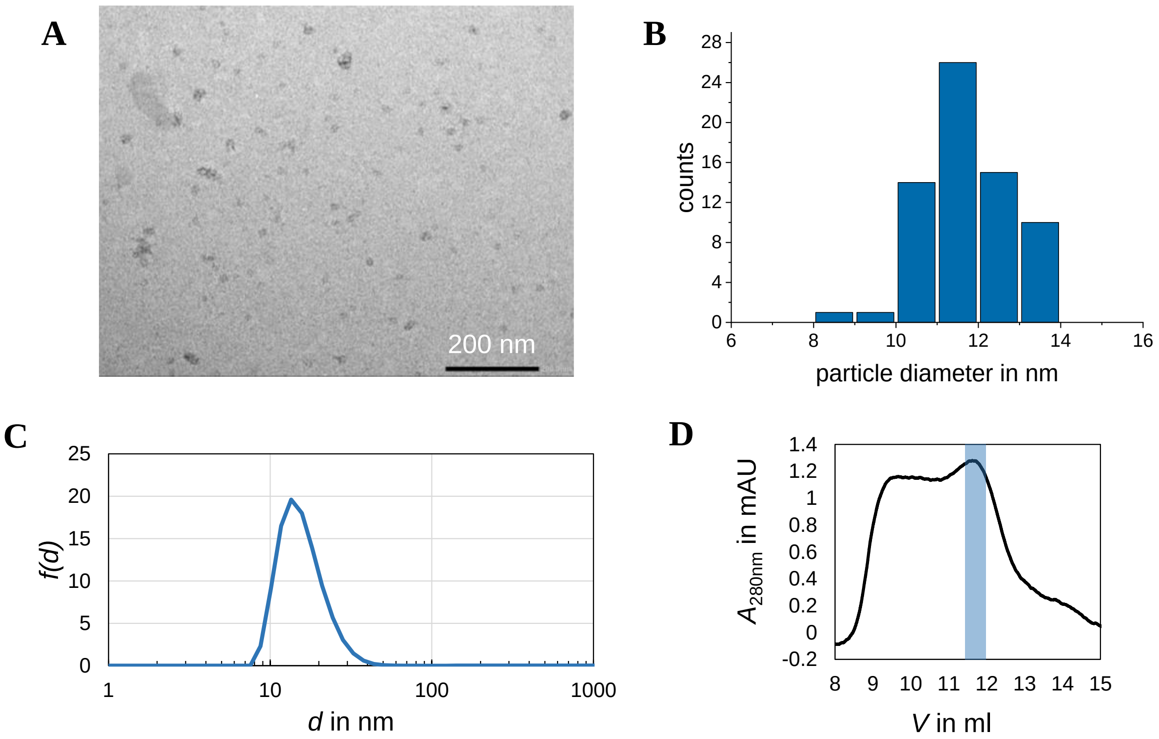

**Figure S3.** Characterization of LpCopA DIBMALPs. **(A)** TEM image of LpCopA DIBMALPs. **(B)** Histogram of size distribution of LpCopA DIBMALPs. Shapiro-Wilk normality test confirmed at the 0.05 level a normally distributed population with a mean of 11.76 nm ± 1.07. **(C)** Volume-weighted particle size distribution *f(d)* with one peak at 13.5 nm. **(D)** Superdex 200 Increase 300/10 GL size-exclusion chromatogram with two major peaks at 9.5 ml and 11.8 ml, which putatively correspond to aggregated particles and LpCopA DIBMALPs.

#### Analysis of background fluorescence of BADAN-labeled single cysteine CopA DIBMALPs

Polymer nanodiscs such as DIBMALPs enable the direct transfer of membrane proteins from native membranes into a discoidal lipid bilayer while preserving their native lipid environment, thereby avoiding transient detergent solubilization. Consequently, an alternative labeling strategy was required for DIBMALPs containing LpCopA mutants that could be incorporated into the DIBMALP preparation protocol. We therefore tested whether BADAN binds specifically to cysteine residues of LpCopA in harvested and resuspended *E. coli* membranes. To reduce nonspecific BADAN binding, the labeled membranes were washed five times to promote partitioning of BADAN into the aqueous phase. During DIBMALP formation, DIBMA was added to disrupt the vesicular bilayer structure, and an equal membrane wet mass of unlabeled *E. coli* membranes lacking overexpressed LpCopA was added to the labeled membranes.

As a control, DIBMALPs were generated from membranes of an *E. coli* strain expressing a cysteine-free LpCopA (cf CopA) mutant, allowing assessment of nonspecific BADAN interactions with *E. coli* lipids. The results are shown in Figure S4. Comparison of the fluorescence intensities at the emission maximum revealed an approximately 50% lower signal for cf CopA. In addition, the emission band shapes of the two variants differed substantially. The emission spectrum of BADAN@C382-LpCopA was red-shifted relative to that of cf CopA, consistent with BADAN covalently bound to a cysteine residue within the transmembrane channel residing in a more hydrophobic environment than BADAN associated with the lipid bilayer or interacting nonspecifically with protein amine groups. Although nonspecific BADAN binding could not be completely eliminated in this batch-labeling approach, a clearly distinguishable signal originating from a single cysteine site was obtained.

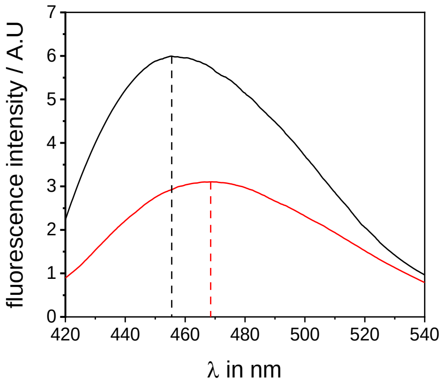

**Figure S4.** Analysis of background fluorescence of BADAN-labeled mutant CopA DIBMALPs. The black spectrum shows the steady state fluorescence spectrum of BADAN@382 CopA, while the red spectrum represents BADAN labeled cf CopA. Emission maxima are indicated by vertical, dashed lines. To evaluate the unspecific BADAN binding between the CopA variants, their fluorescence intensity has been compared normalized to the same protein concentration.

#### Probing site-specific fluorescence shifts induced by increasing osmotic stress of various membrane mimetics

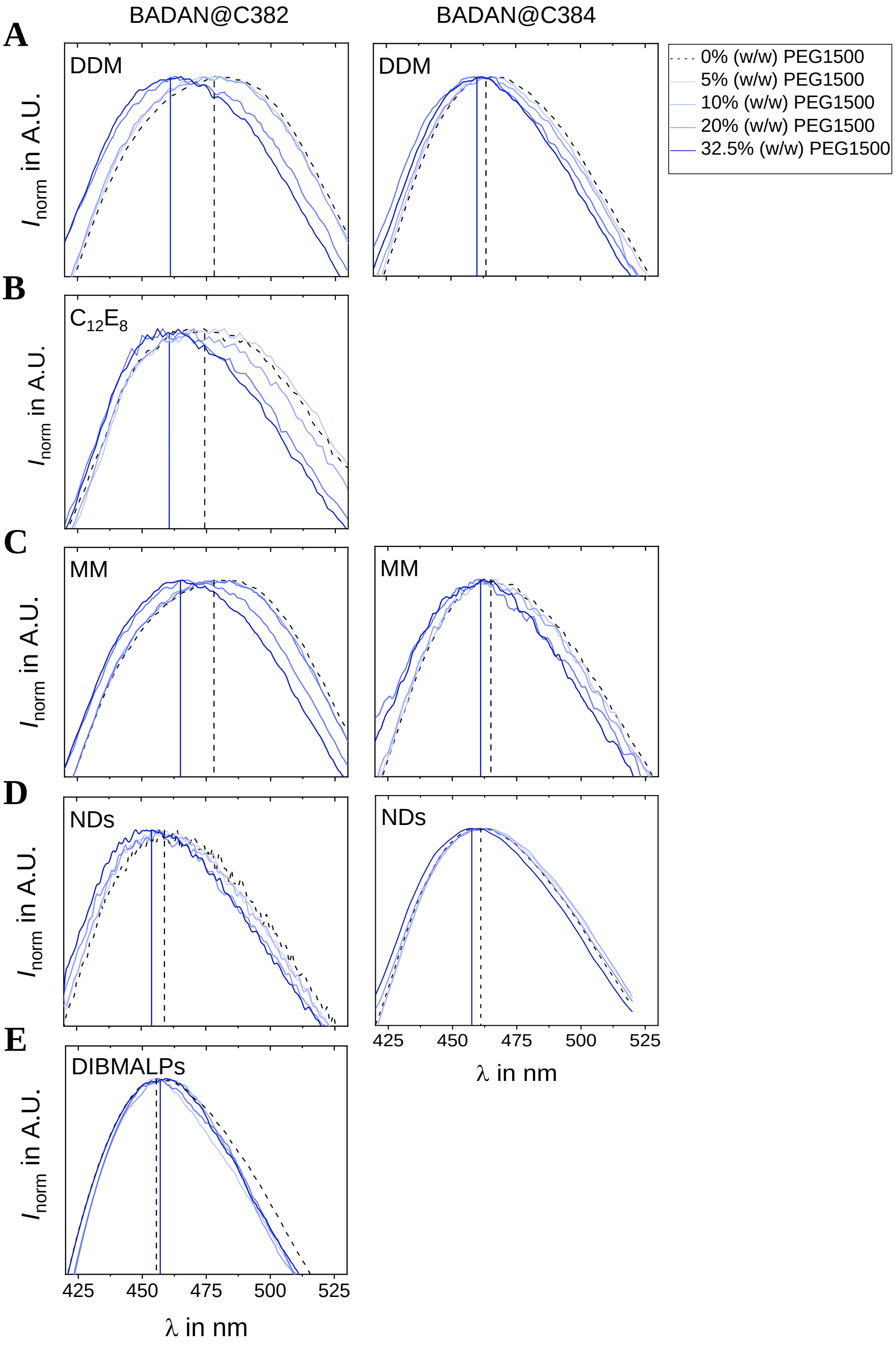

**Figure S5.** Steady state fluorescence spectra of BADAN-labeled single-cysteine LpCopA mutants in different membrane mimetic systems, probed with increasing concentrations of PEG-1500. Left-handed site windows correspond to the fluorescence output of the single cysteine mutant with BADAN at cysteine 382, whereas the right-handed represent mutant with BADAN at cysteine 384. Each row shows another membrane mimetic system in which the protein mutants have been incorporated **(A)** DDM detergent micelles, **(B)** C12E8 detergent micelles, **(C)** Mixed micelles, **(D)** MSP1E3D1 nanodiscs and **(E)** DIBMALPs. The color bar code with the corresponding PEG-1500 concentrations can be found on the upper right site.

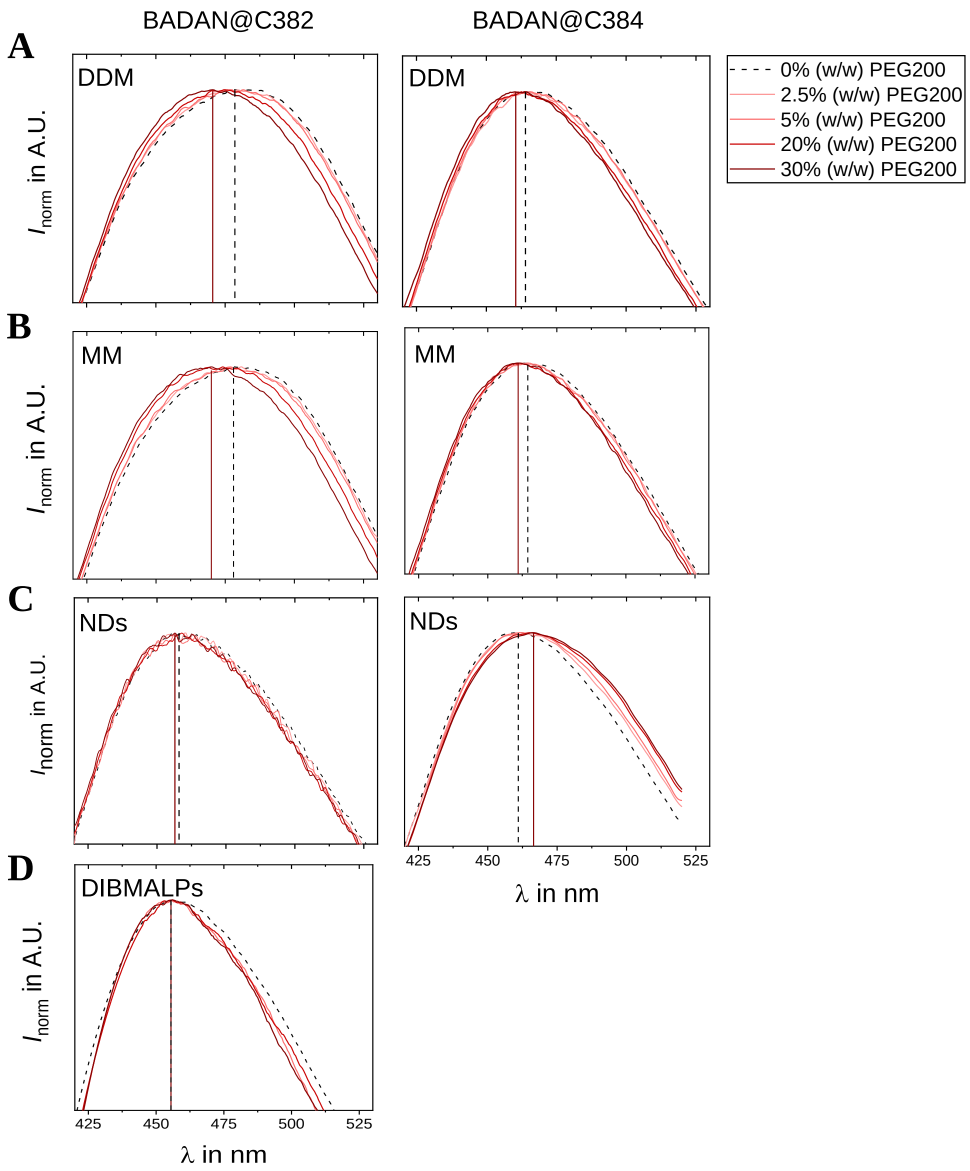

**Figure S6.** Steady state fluorescence spectra of BADAN-labeled single-cysteine LpCopA mutants in different membrane mimetic systems, probed with increasing concentrations of PEG-200. Left-handed site windows correspond to the fluorescence output of the single cysteine mutant with BADAN at cysteine 382, whereas the right-handed represent mutant with BADAN at cysteine 384. Each row shows another membrane mimetic system in which the protein mutants have been incorporated **(A)** DDM detergent micelles, **(B)** Mixed micelles, **(C)** MSP1E3D1 nanodiscs and **(D)** DIBMALPs. The color bar code with the corresponding PEG200 concentrations can be found on the upper right site.
